## Supplementary figures and images for "KLC4 shapes axon arbors during development and mediates adult behavior"

### Figure Supplement 1

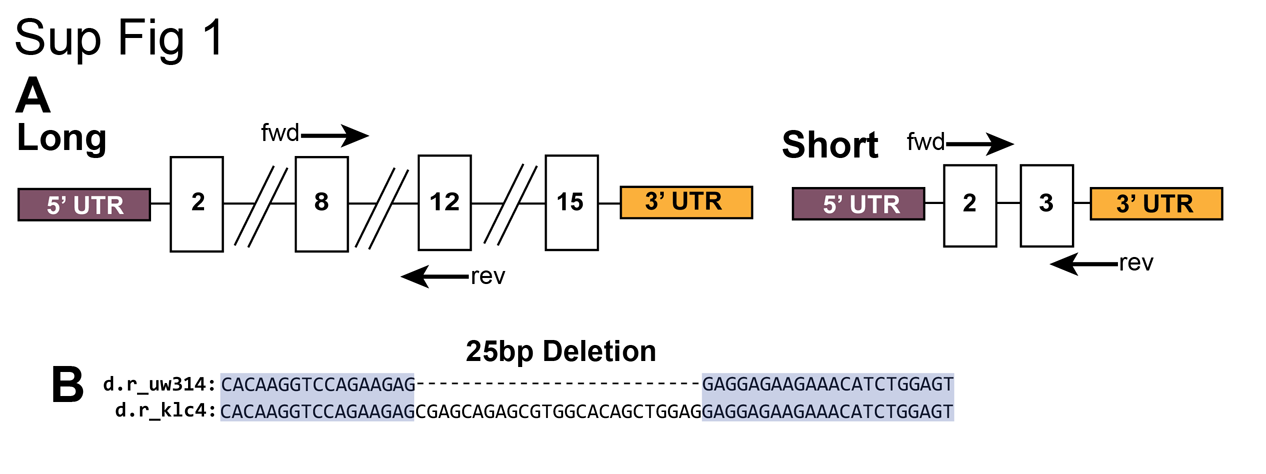

### Figure Supplement 2

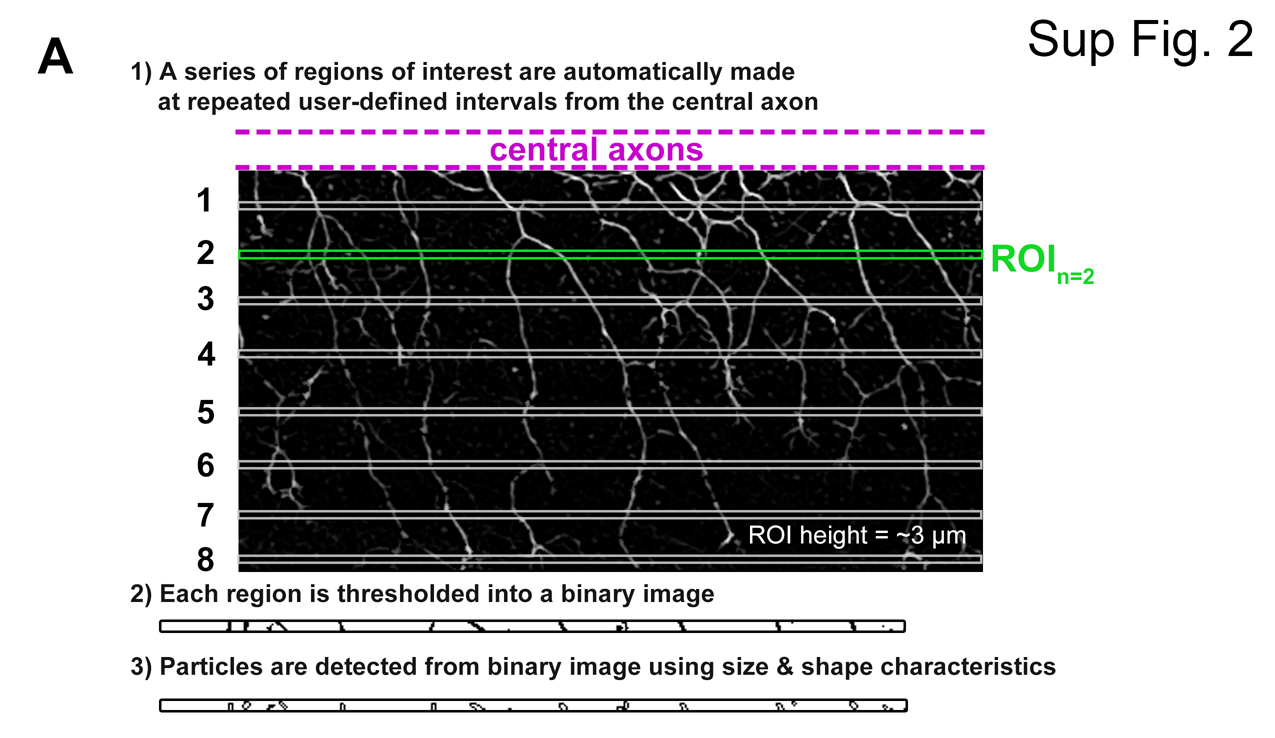

### Figure Supplement 4

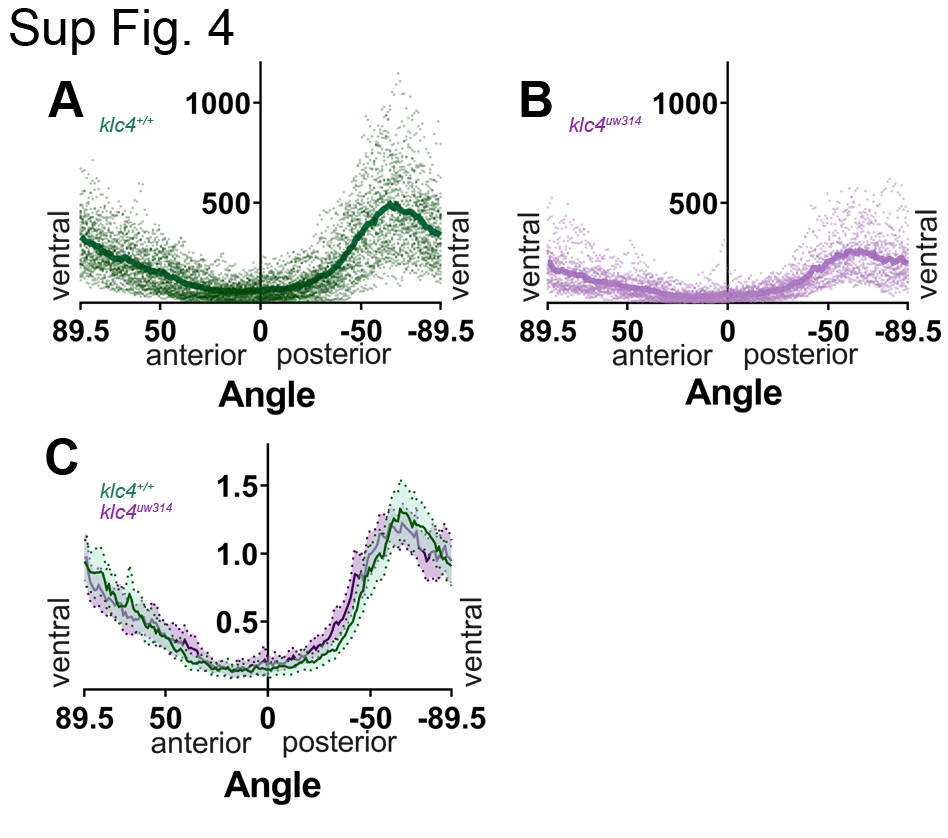

### Figure Supplement 8

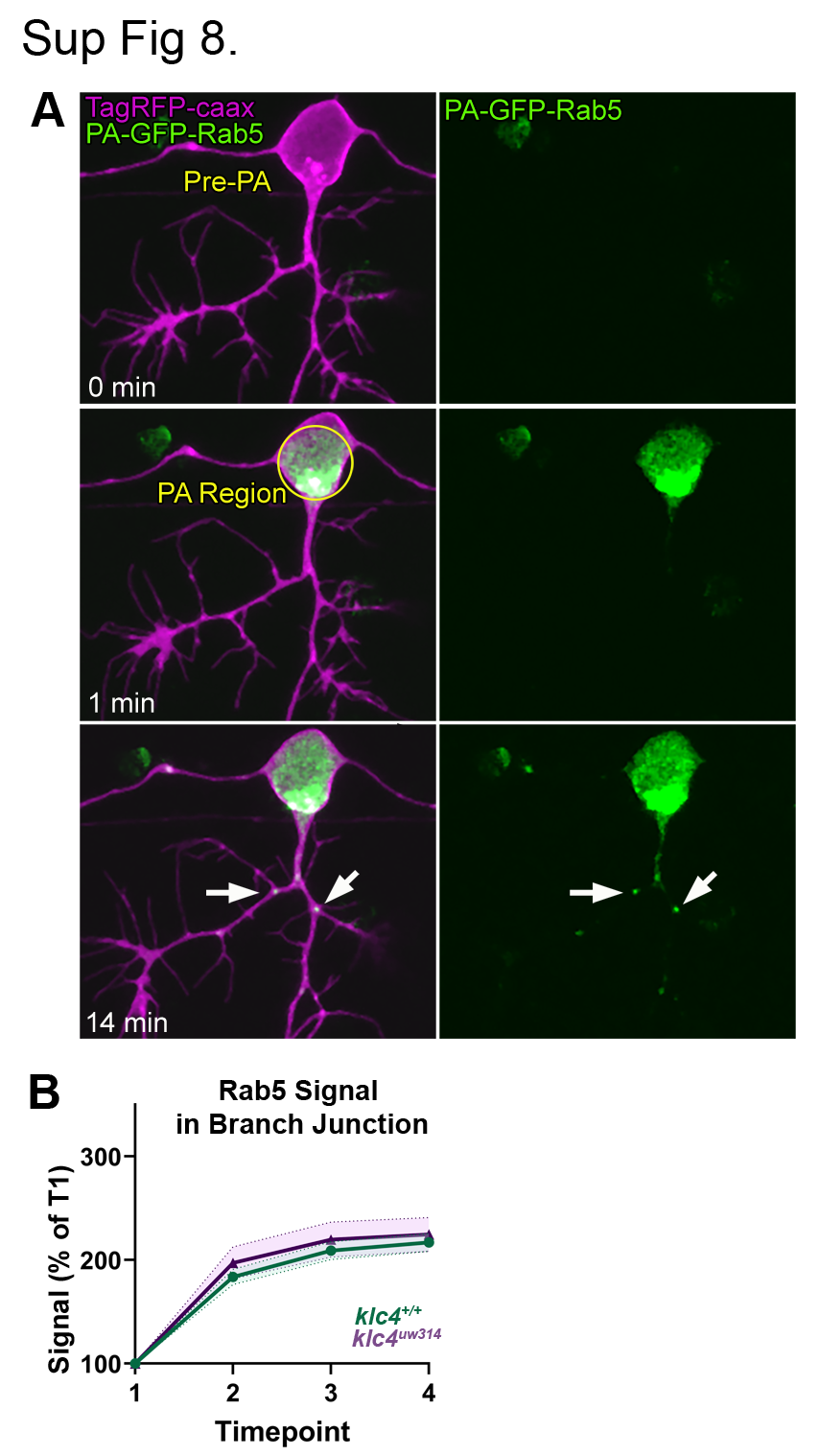

### Figure Supplement 9

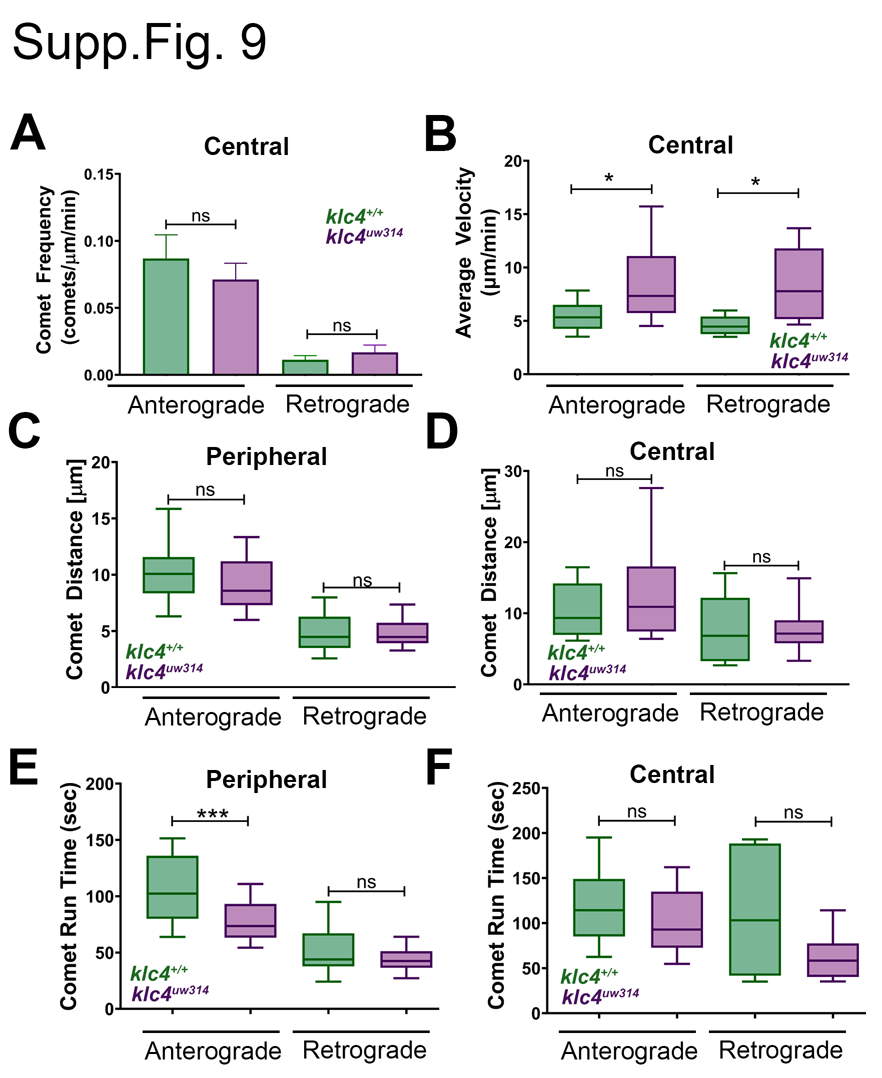

### Figure Supplement 11

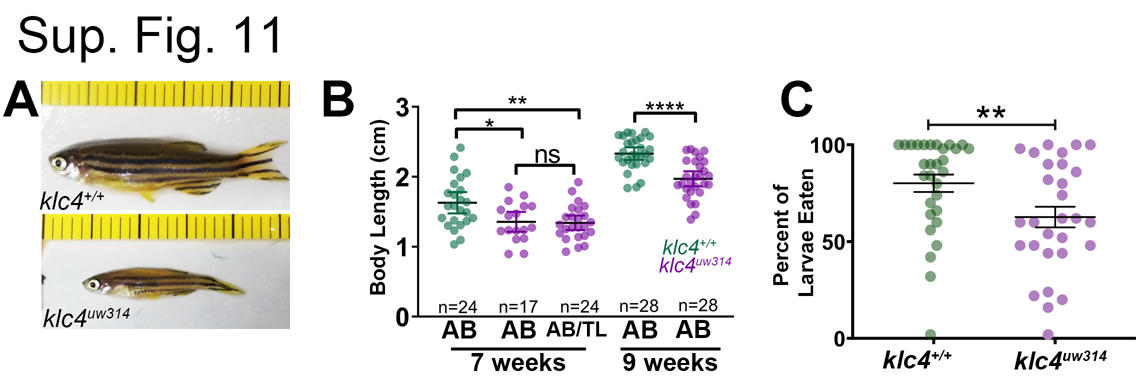

### Figure Supplement 12

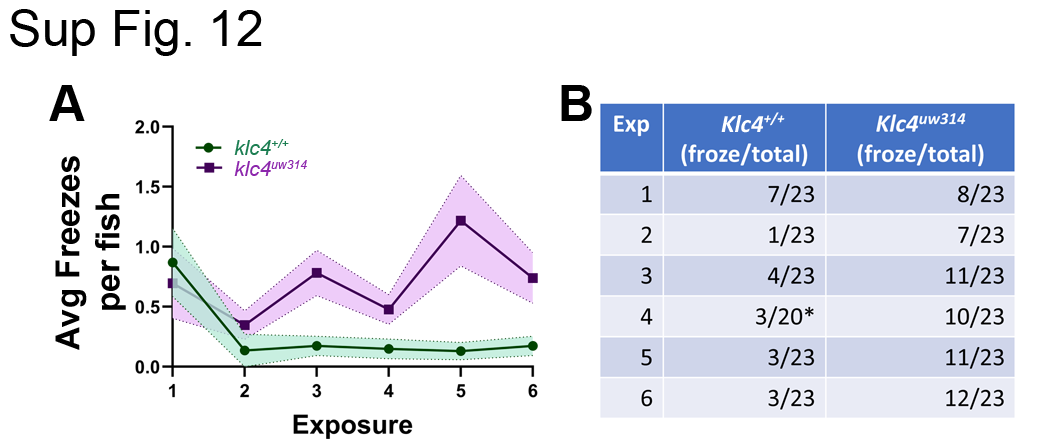
